# Revealing microbial assemblage structure in the human gut microbiome using latent Dirichlet allocation

**DOI:** 10.1101/664219

**Authors:** Shion Hosoda, Suguru Nishijima, Tsukasa Fukunaga, Masahira Hattori, Michiaki Hamada

## Abstract

Recent research has revealed that there are various microbial species in the human gut microbiome. To clarify the structure of the human gut microbiome, many data mining methods have been applied to microbial composition data. Cluster analysis, one of the key data mining methods that have been used in human gut microbiome research, can classify the human gut microbiome into three clusters, called enterotypes. The human gut microbiome has been suggested to be composed of the microbial assemblages or groups of co-occurring microbes, and one human gut microbiome can contain several microbial assemblages. However, cluster analysis can cluster samples into groups without capturing minor assemblages. In addition, a reliable method of assemblage detection has not been established, and little is known about the distributions of microbial assemblages at a population-level scale. Accordingly, the purpose of this study was to clarify the microbial assemblages in the human gut microbiome. In this study, we detected gut microbiome assemblages using a latent Dirichlet allocation (LDA) method, which was first proposed for the classification of documents in natural language processing. We applied LDA to a large-scale human gut metagenome dataset and found that a four-assemblage LDA model can represent relationships between enterotypes and assemblages with high interpretability. This model indicates that each individual tends to have several assemblages, and each of three assemblages corresponded to each enterotype. However, the C-assemblage can exist in all enterotypes. Interestingly, the dominant genera of the C-assemblage (*Clostridium, Eubacterium, Faecalibacterium, Roseburia, Coprococcus*, and *Butyrivibrio*) included butyrate-producing species such as *Faecalibacterium prausnitzii*. Finally, we revealed that genera mainly appearing in the same assemblage were correlated to each other. We conducted an assemblage analysis on a large-scale human gut metagenome dataset using LDA, a powerful method for detection of microbial assemblages. This approach has the potential to reveal the structure of the human gut microbiome.

## Introduction

The human gut microbiome varies greatly from person to person, depending on differences among human populations [1] and dietary habits [2]. The differences in gut microbial compositions affect host health and physiology [3], and in some cases, altered microbial compositions are associated with diseases such as inflammatory bowel disease [4], type-1 diabetes [5], colorectal cancer [6], and autism [7, 8]. Recent development of metagenome sequencing technologies have enabled investigations of gut microbial compositions of individuals with ease and rapidity, and many large-scale research projects focused on the human gut microbiome have been conducted [1, 9, 10, 11]. At present, by applying various data mining methods to these massive metagenomic datasets, the structure of the human gut microbiome and the relationship between a host’s phenotype and its gut microbial profile can be revealed.

Cluster analysis of samples is one of the widely used data mining methods in metagenomic research. In this approach, individuals are clustered into groups based on similarities in their microbial profiles. For example, Arumugam *et al*. discovered that the gut microbial profiles of individuals can be classified into three types called enterotypes using the partitioning around medoids (PAM) clustering method [12]. In another example, Ding and Schloss reported that the human gut microbiome has considerable inter-individual variation, but the cluster type of an individual was almost unchanged in the sampling period, using the Dirichlet multinomial mixture (DMM) clustering method [13, 14]. Although cluster analysis is a powerful approach for revealing the overall structure of human gut microbiomes, this analysis is strongly affected by the dominant microbes in each individual. Therefore, cluster analyses of samples may ignore the existence of non-dominant but shared microbes among individuals (Fig. 1).

**Figure 1:**
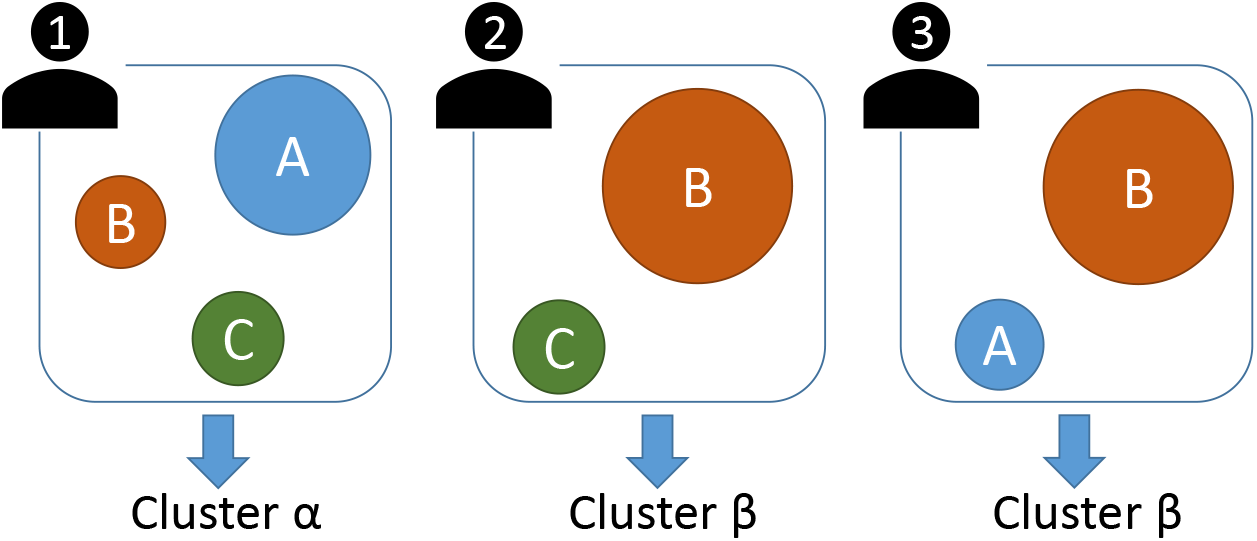
Schematic illustration of microbial assemblage and cluster analysis for the human gut microbiome, where A, B, and C show microbial assemblages with circle size indicating abundance. The cluster of each individual was determined by the dominant assemblage. However, a cluster analysis cannot capture the non-dominant but shared microbes among samples like those comprising assemblage C.

An alternative data mining method is microbial assemblage analysis, which clusters microbes into some assemblages. Here, following Boon *et al*. [15], we define assemblages as groups of microbes that are expected to co-occur. In this view, an individual can have several microbial assemblages. Therefore, this analysis can capture assemblages consisting of non-dominant microbes, unlike a cluster analysis of samples. Shafiei *et al*. developed BioMiCo, which is a Bayesian probabilistic model for microbial assemblage analysis, and discovered host-specific assemblages in human gut metagenomic time-series data [16]. Recently, Cai *et al*. also explored microbial assemblages using non-negative matrix factorization methods and identified a shift of microbial assemblages in one individual [17]. Microbial assemblage analysis has also been used to track sources of contamination in metagenomic research [18] and to detect assemblage-level metabolic interactions [19], but microbial assemblage analysis based on a massive metagenome dataset has not yet been conducted. As such, the large-scale assemblage structure of human gut microbiomes and the relationship between microbial assemblages and enterotypes are still unknown.

In this study, we conducted a microbial assemblage analysis of a large-scale human gut metagenomic dataset in order to reveal the structures of microbial assemblages in the human gut microbiome. To detect assemblages, we used the latent Dirichlet allocation (LDA) method, which is an unsupervised probabilistic model [20]; LDA was first proposed for the classification of documents in natural language processing, and this method is now widely used in bioinformatics fields such as transcriptome analysis [21], pharmacology [22], and gene function prediction [23]. LDA allows one microbe to be assigned to multiple clusters; this characteristic serves as an advantage for modeling microbiome clusters because dependency among microbes result from metabolic functions shared by several microbes, and microbes are interchangeable with other microbes having the same metabolic functions. Yan *et al*. developed MetaTopics [24] and applied it to the human oral metagenomic dataset and the human gut metagenomic dataset. However, they applied MetaTopics to small datasets including fewer than 200 samples and have neither performed detailed analyses nor discussed their findings in detail.

In the present study, we first considered the number of microbial assemblages based on the relationships between microbial assemblages and enterotypes. Next, we found that a four-assemblage model has high interpretability in the context of a large-scale human gut microbiome dataset and discovered that an individual may have not just one microbial assemblage but several assemblages in many cases. Then, we investigated the relationships between enterotypes and microbial assemblages and revealed that three assemblages correspond to each enterotype but that one assemblage can exist in all enterotypes. Thereafter, we examined the human population-level differences in microbial assemblages and found the existence of population-independent microbial assemblages. Finally, we estimated the functions of each assemblage by applying LDA to the functional profiles that with the same samples as the genus data. They are referred to as “functional assemblages” in later analyses.

## Materials and methods

### Metagenomic dataset and preprocessing methods

We used the large-scale human gut metagenome dataset constructed by Nishijima *et al*. [25]. This dataset consisted of gut metagenomic data from 861 healthy adults from 12 countries. The taxon of each sequencing read was assigned by mapping the read to a reference genome dataset consisting of 6,149 microbial genomes.

We used genus as the taxonomic rank for each sequencing read because genus rank has been used in previous studies including enterotype analysis. We calculated the normalized number of occurrences of each genus in each individual so that the total number of occurrences of all genera per individual would be 10,000 (because LDA cannot be applied to datasets consisting of fractional values). After these preprocessing steps, the number of different genera included in the dataset became 252.

In functional assemblage analysis, we used the Kyoto Encyclopedia of Genes and Genomes (KEGG) [26] orthology annotated data as functional profiles. This dataset was also constructed by Nishijima *et al*. [25]. We calculated the normalized number of occurrences of each KEGG orthology in each individual, such that the total number of occurrences of all genera per individual would be 100,000. After these preprocessing steps, the number of different KEGG orthologies included in the dataset was 1790.

### LDA for modeling the human gut microbiome

The probabilistic LDA model [20] can be used to estimate *K* microbial assemblages in whole human gut metagenomic datasets, where *K* is a given parameter. In the LDA model, each metagenome sample *s_i_* (*i* ∈ {1, …, 861}) has a multinomial distribution with parameter 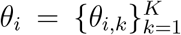 over microbial assemblages where *θ_i,k_* is the occurrence probability of the *k*-th assemblage in the *i*-th sample; each microbial assemblage *a_k_* (*k* ∈ {1, …, *K*}) has a multinomial distribution with parameter 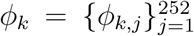 over genera *g_j_* (*j* ∈ {1, …, 252}), where *ϕ_k,j_* is the occurrence probability of the *j*-th microbe in the *k*-th assemblage.

A microbial assemblage with high probability in an individual means that the individual tends to have that particular microbial assemblage in the gut microbiome, and a genus with high probability in a microbial assemblage means that the microbial assemblage tends to have that particular genus. In addition, the LDA model has prior distributions of *θ_i_* and *ϕ_k_* provided by the Dirichlet distribution whose hyperparameter is *α* and *β*, respectively. In this research, we used 0.1 and 0.05 as the initial values of all the elements of *α* and *β*, respectively.

The LDA parameters (*θ* and *ϕ*) can be learned from the dataset in an unsupervised manner. Various parameter inference methods for the LDA model have been proposed, and we used the variational Bayes (VB) method [20], which maximizes the approximation of a marginal likelihood, called the variational lower bound (VLB) score, by updating the parameter iteratively from random initial parameters. We concluded the iteration of the parameter update when the change in the VLB score between the previous and current step was less than 10^−6^. Finally, we estimated each *θ_i_* and *ϕ_k_* as the expectation values of the distribution estimated by the VB method. We conducted 10 trials for each *K* = 2, 3, 4, 5 and adopted the estimated parameter with the highest VLB score among all trials for each *K*. In addition, we updated the hyperparameters *α* and *β* from the initial values using a fixed point iteration method in the parameter learning step [27]. Based on previous research about LDA hyperparameter settings [28], we estimated the parameter so that each element of *α* differed from the others but all elements of *β* have the same value. Furthermore, in the functional assemblage analysis, the experimental conditions and methods were the same as those explained above. In addition, *K* was set at the same number as the microbial assemblages.

### Calculation method of entropy scores

To quantify the bias in the occurrence frequency for each element in the estimated probability distribution, we calculated the entropy scores of the occurrence distribution of the assemblage for each sample and for each genus. In a multinomial distribution, a high entropy score means that the distribution is similar to the uniform distribution, and a low score means that the distribution tends to take a specific value. The entropy score *H*(*a*|*s_i_*) of the assemblages 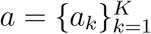 for each sample *s_i_* was calculated as follows:

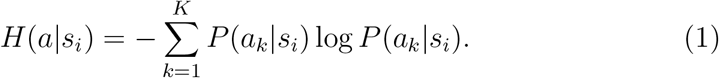

As *P*(*a_k_*|*s_i_*) is equal to *θ_i,k_*, we can directly calculate this score using the estimated LDA parameters. The entropy score *H*(*a*|*g_i_*) of the assemblages 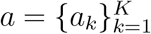 for each genus *g_j_* was calculated as follows:

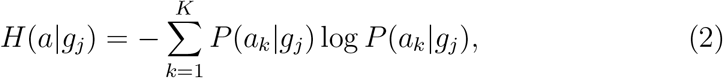

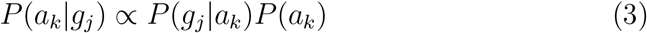

where *P*(*g_j_*|*a_k_*) is equal to *ϕ_k,j_* and the average probability of all *θ_i,k_* was used as *P*(*a_k_*).

## Results

### Cluster analysis of the human gut microbiome enterotypes

In order to investigate the relationship between enterotypes and assemblages in the following analysis, we classified individual samples into three clusters using the PAM clustering method. We verified that the dominant genus in each identified cluster were *Bacteroides, Prevotella*, and *Blautia*, and these genera were specific to each cluster (Additional File 1, Figure S1). These results are consistent with the previous enterotype research, in which three enterotypes were identified in the human gut microbiome: *Bacteroides* dominant type, *Prevotella* dominant type, and *Ruminococcus* and *Blautia* dominant type [12]. Hence, we called these clusters the B-type, P-type, and R-type, respectively.

### Analysis of the human gut microbial assemblage profiles estimated by LDA

We conducted estimates of the 2–5-assemblage LDA model parameters to find the model that has the highest interpretability of relationships between enterotypes and assemblages. Fig. 2 shows the assemblage distributions for each enterotype obtained by each model. The two-assemblage model identified the B-type specific assemblage and P- and R-type specific assemblage (IDs 1 and 2 in Fig. 2a). The three-assemblage model estimated the assemblages corresponding to each enterotype (Fig. 2b). In addition to these enterotype-specific assemblages, the four- and five-assemblage models estimated the general assemblage that appears in *all* the enterotypes (Fig. 2cd). The strength of LDA is that it is possible to obtain such an assemblage. The five-assemblage model estimated two general assemblages (IDs 4 and 5 in Fig. 2d), and it seems to be difficult to identify these assemblages because they are equivalent to enterotypes. In other words, the two assemblages of the five-assemblage model had the same occurrence pattern for enterotypes. Therefore, we used the four-assemblage model in this study. Note that the existence of a general assemblage is not trivial in models with four or more assemblages because there are not always genera that appear in all enterotypes. In the following analysis, we call the assemblages with IDs 1, 2, and 3 the “B-assemblage,” “P-assemblage,” and “R-assemblage,” respectively, because these assemblages appeared specifically in the B-, P-, and R-type individuals, respectively (Fig. 2c). In addition, we call the assemblage with ID 4 the “C-assemblage.”

**Figure 2:**
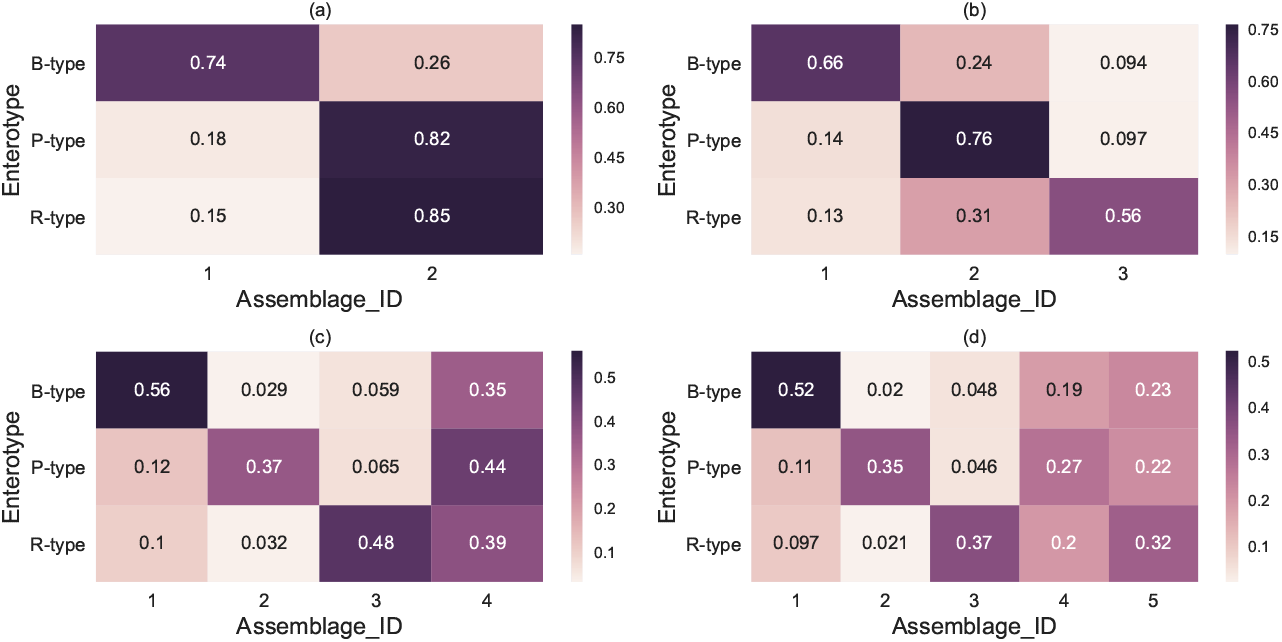
Assemblage distributions for each enterotype. Each row shows a distribution obtained by averaging the estimated assemblage distributions of individuals corresponding to each enterotype. The *x*- and *y*-axes represent the microbial assemblages and enterotypes, respectively. Darker colors indicate higher probabilities, and each number inside the partition indicates a different probability. (a), (b), (c), and (d) indicate 2–5-assemblage LDA models, respectively.

Next, we investigated the kinds of genera that constituted each microbial assemblage. Fig. 3 shows the genus distribution of each microbial assemblage estimated by LDA (i.e., 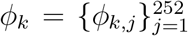 in the previous section). B- and P-assemblages mainly consisted of one dominant genus, *Bacteroides* and *Prevotella*, and the relative frequencies were 71% and 66%, respectively. On the other hand, R- and C-assemblages consisted of genera with medium occurrence frequencies. The genera that constituted the R-assemblage were *Blautia* (22%), *Bifidobacterium* (20 %), and *Ruminococcus* (8.6 %), among others. The C-assemblage consisted of *Clostridium* (18%), *Eubacterium* (15 %), and unclassified *Firmicutes* (13 %), among others.

**Figure 3:**
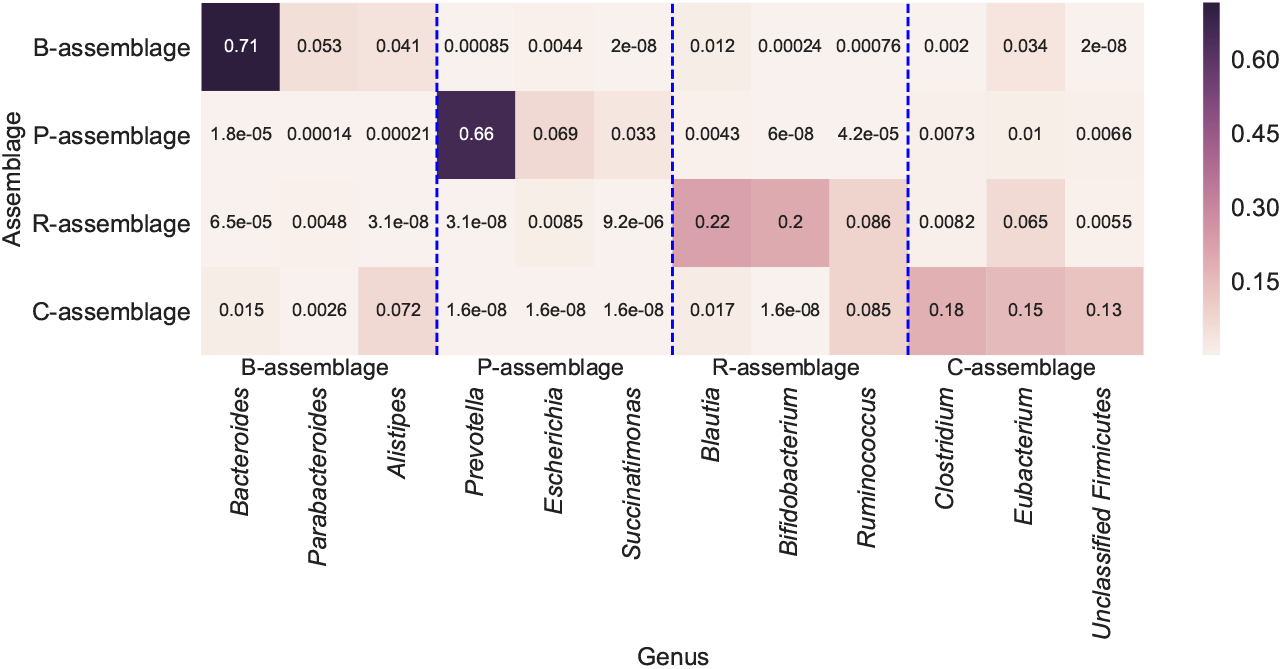
The estimated genus distribution of each microbial assemblage. The *x*- and *y*-axes represent genera and assemblages, respectively. We displayed only the three genera with the highest probability in each assemblage.

In the LDA model, a genus can appear in several microbial assemblages. We investigated whether genera occurred in just one specific assemblage or not using the entropy scores of the assemblage occurrence distributions for genera (Eq. 2). Fig. 4a shows a histogram of the entropy scores for all genera, and two peaks, at 0.00–0.125 and 0.50–0.75, were observed in the distribution. The former peak represents assemblage-specific genera, and *Bacteroides* and *Prevotella* belonged to this group (Additional File 1, Table S1). The latter peak represents a genus appearing in several but not all assemblages, and *Ruminococcus* and *Blautia* belong to this group (Additional File 1, Table S1). Several genera showed high entropy scores, which indicates they are universal genera among assemblages (Additional File 1, Figure S2). As such, occurrence tendencies of the assemblages for each genus varied (Additional File 1, Table S1).

**Figure 4:**
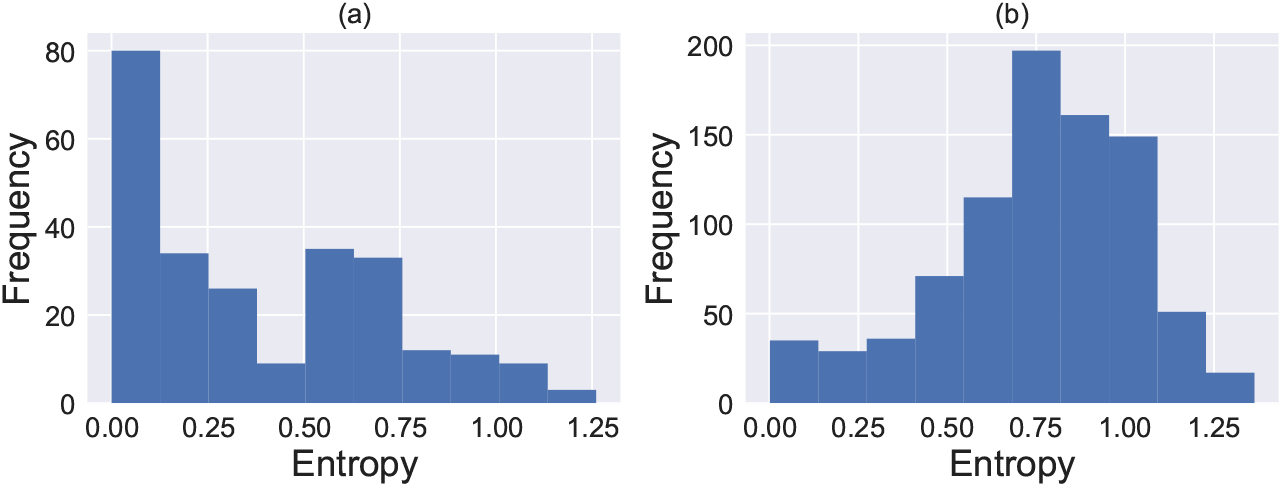
Histograms of the entropy scores of the microbial assemblage (a) for each genus (Eq. 2) and (b) for each individual (Eq. 2). The *x*- and *y*-axes represent entropy and the number of samples, respectively.

Then, we calculated the entropy scores of the assemblage occurrence distributions (Eq. 1) for each individual (Fig. 4b). The distribution of the entropy score was unimodal, and the median was 0.7805. These results suggest that most individuals have multiple but not all microbial assemblages. In addition, we examined compositions of microbial assemblages for each individual (Additional File 1, Figure S3) and found that co-occurrence tendencies between microbial assemblages were not uniform. That is, P- and R-assemblages tended to exist exclusively in each sample, but B- and C-assemblages could coexist with other assemblages.

### Relationships between microbial assemblages and enterotypes

We investigated the relationships between microbial assemblages and enterotypes to reveal how assemblages appear in enterotypes.

Fig. 5a shows the relationship between microbial assemblages and enterotypes. While B-, P-, and R-assemblages correspond to three enterotypes as mentioned above, the C-assemblage was observed in all three enterotypes. This result suggests that the dominant microbial assemblages of each enterotype differ from each other but there is an assemblage of non-dominant genera that can exist in all enterotypes. To confirm this interpretation, we investigated the occurrence tendency of the genera mainly appearing in each assemblage. Here, each genus was regarded as mainly appearing in the assemblage of the highest *P*(*a_k_*|*g_i_*) (Eq. 3). Fig. 6 shows the relative abundance of genera in each individual, which was constructed by Nishijima *et al*. [25]. We found that the genera mainly appearing in the C-assemblage occurred in all enterotypes and that the genera mainly appearing in the R-assemblage occurred in R-type individuals. These results were supported by the estimated parameters shown in Fig. 5a. In addition, the genera mainly appearing in the B-assemblage, other than *Bacteroides* occurring in the B-type, was 2.05 times as frequent as in the other enterotypes. The genera mainly appearing in the P-assemblage, other than *Prevotella* occurring in P-type, was 3.00 times as frequent as in the other enterotypes.

**Figure 5:**
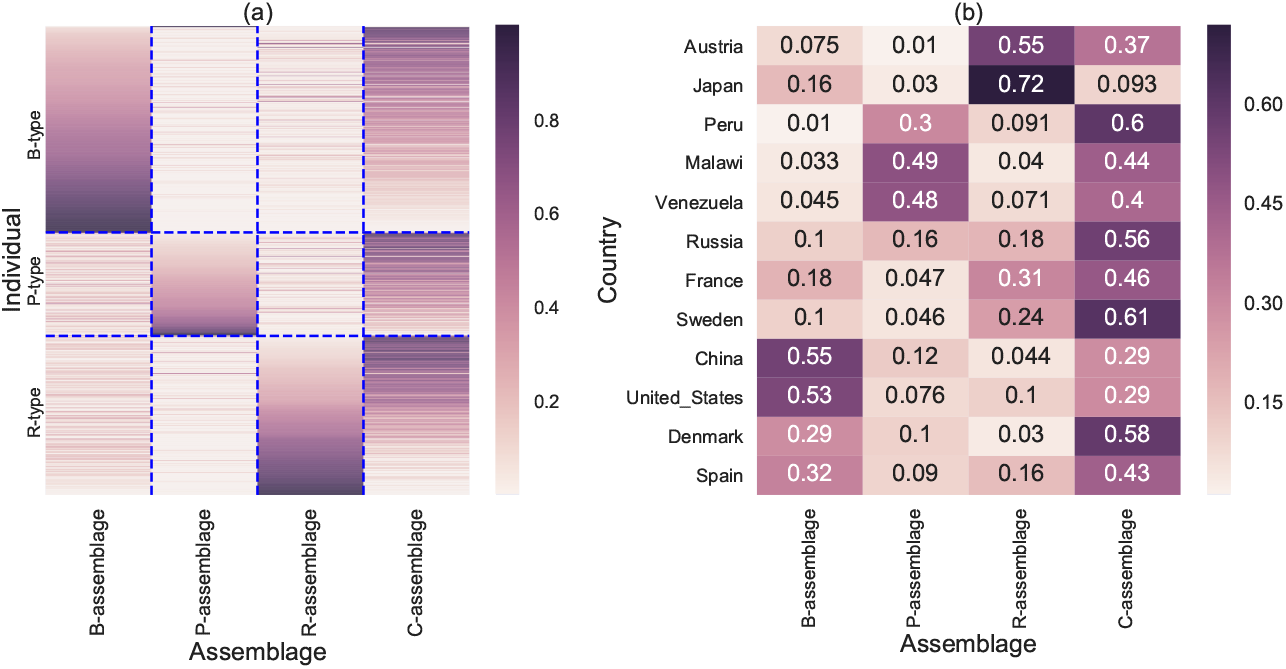
(a) Estimated microbial assemblage distribution for each individual. The *x*- and *y*-axes represent microbial assemblages and individual samples, respectively. Individuals are segregated by the enterotype and sorted by the B-assemblage, the P-assemblage, and the R-assemblage, respectively. (b) Average assemblage distributions for each country. Each row shows a distribution obtained by averaging the estimated assemblage distributions of individuals corresponding to each country. The *x*- and *y*-axis represent the microbial assemblage and the country of the individual, respectively.

**Figure 6:**
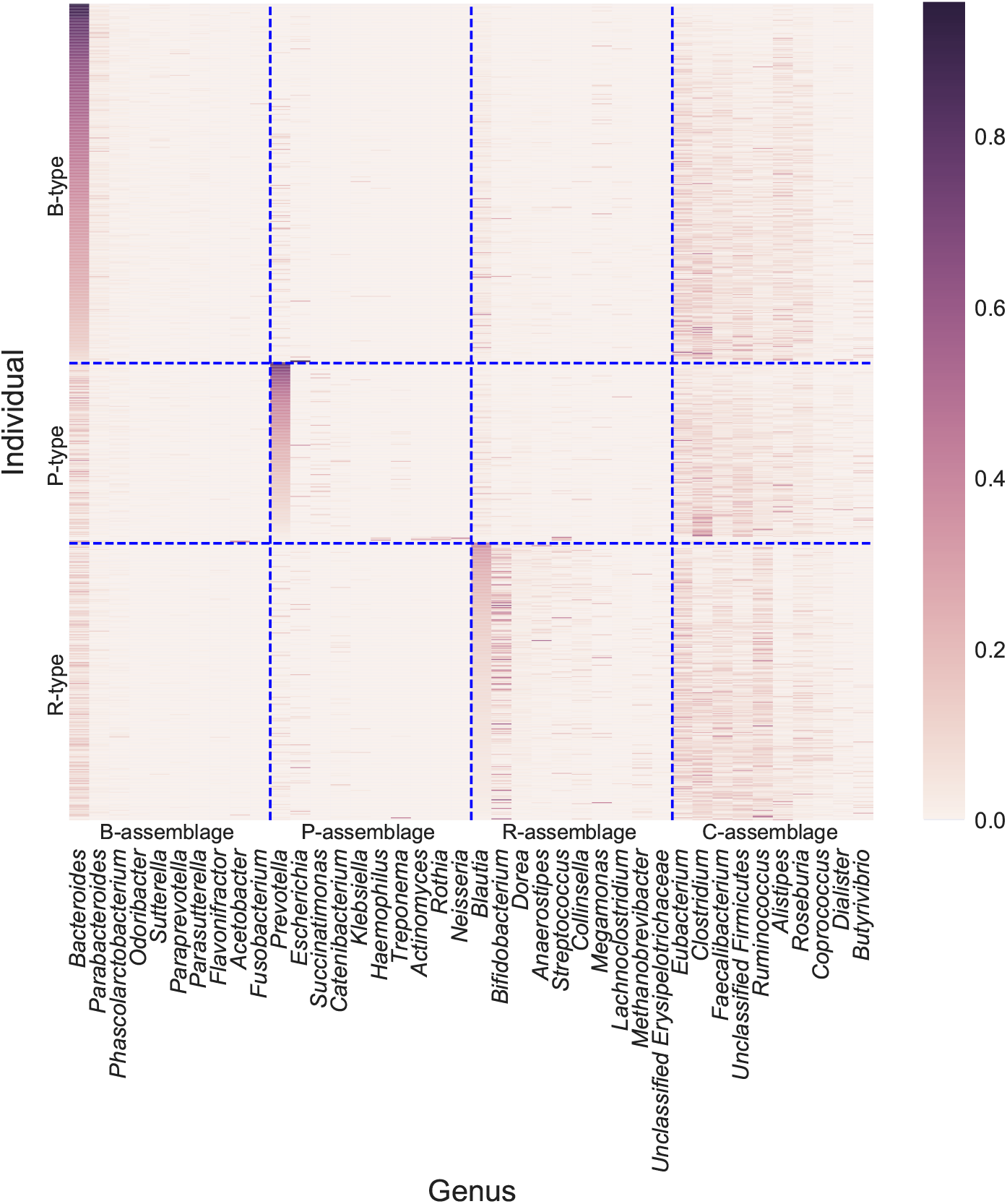
Relative abundance of genera in each individual. The *x*- and *y*-axes represent the genus and individuals, respectively. Individuals are divided by the enterotype and sorted by *Bacteroides, Prevotella*, and *Blautia*, respectively. Genera are divided by the assemblage that they mainly appear in and are sorted by the abundance of each genus. Each genus was regarded as mainly appearing in the assemblage of the highest *P*(*a_k_*|*g_i_*), where *a_k_* and *g_i_* are the *k*-th assemblage and the *i*-th genus, respectively (Eq. 3 in the main text).

### Correlations between microbes in the same or different assemblages

We examined the relationship between microbes in the same or different assemblages to determine the significance of the assemblages. Fig. 7a shows the Spearman’s correlation coefficients between the major genera of the B-, P-, R-, and C-assemblages, and suggests that the genera of the same assemblage are correlated with each other. To verify this suggestion, we conducted a correlation test, and most genera from the same assemblage were significantly correlated (Fig. 7b) at *p* < 0.01 (two-sided test, after Benjamini–Hochberg correction[29]). Some genera (*i.e., Eubacterium, Faecalibacterium, Dorea, Ruminococcus*, *Streptococcus*, and *Catenibacterium*) were significantly correlated with many genera in other assemblages. These results are consistent with the fact that their *P*(*a_k_*|*g_i_*) (in Eq.3) is high for multiple assemblages. For example, *Ruminococcus* has a positive correlation with the genera mainly appearing in the R-assemblage. Indeed, *Ruminococcus* has a high association with the R-assemblage even though its main assemblage is the C-assemblage (Additional File 1, Figure S4). These results indicate that the LDA model can capture the assemblages as groups of correlated genera. Fig. 7a also shows high correlation coefficients between the genera mainly appearing in the R-assemblage. This observation may be affected by many Japanese samples, which have high R-assemblage abundance as mentioned below.

**Figure 7:**
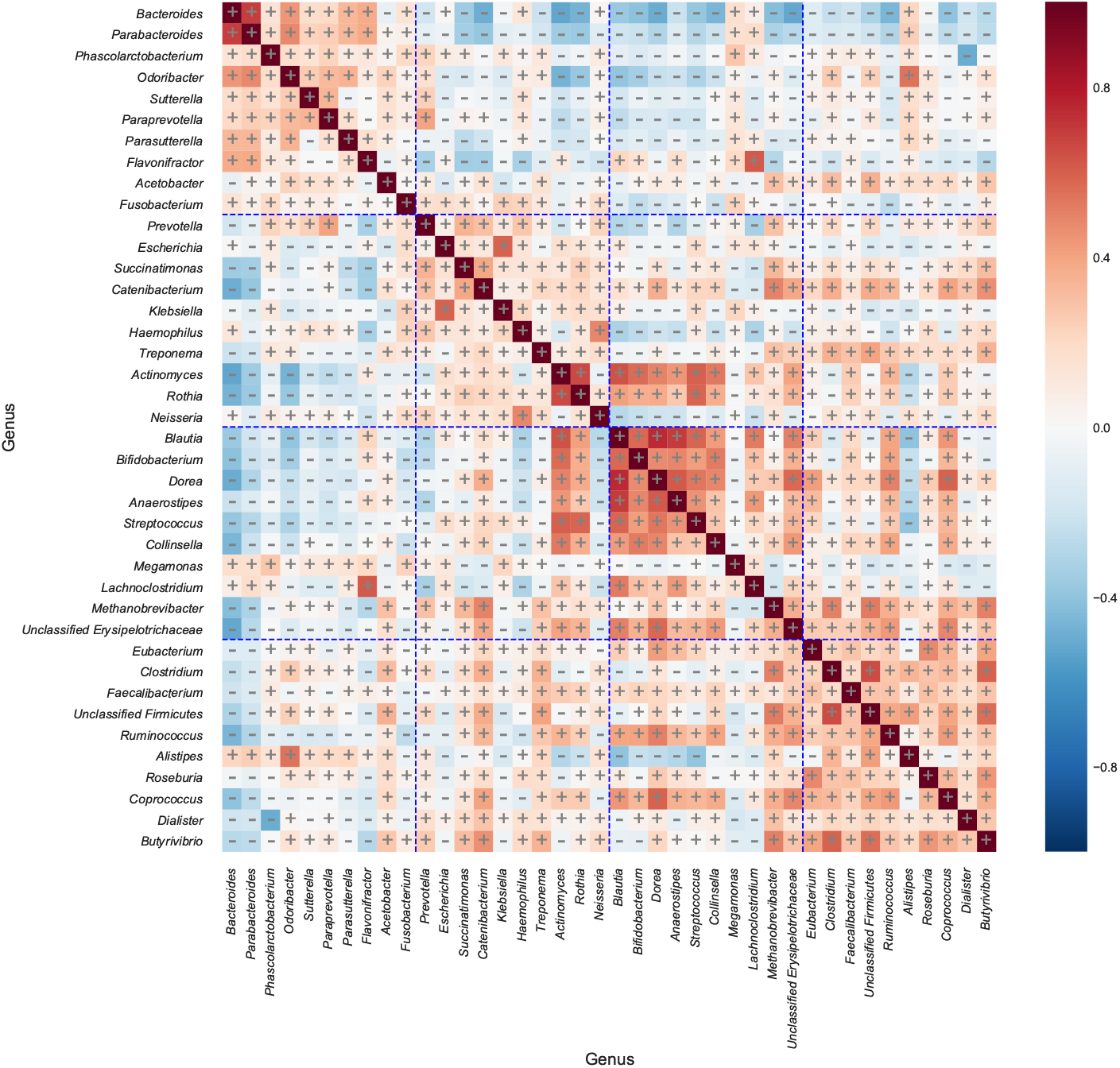
The Spearman’s correlation coefficients among the 20 genera that are major in each enterotype. Both the *x*- and *y*-axes represent genera, which are divided and sorted in the same way as in Fig. 6. Plus and minus signs indicate significant positive and negative correlations, respectively. Significance was determined at *p* < 0.01 (two-sided test, after Benjamini–Hochberg correction).

### Relationships between microbial assemblages and countries

We investigated the relationships between microbial assemblages and countries to observe tendencies of the assemblages among host countries.

Fig. 5b shows the average assemblage distributions of individuals for each country. We discovered that the occurrence distributions of microbial assemblages vary from country to county; for example, Japan and Austria tend to have R-assemblages while Malawi and Venezuela tend to have P-assemblages. On the other hand, the C-assemblage was frequently found in all countries except Japan. Note that Nishijima *et al*. reported that the Japanese gut microbiome was characterized by low abundance of *Clostridium* and unclassified *Firmicutes*, which are main components of the C-assemblage (Table 2) based on the same dataset [25]. Incidentally, *Eubacterium* and *Faecalibacterium*, which are abundant genera in the C-assemblage, were not less abundant in the Japanese population in comparison with other countries (Additional File 1: Figure S5).

### Correlations between microbial assemblages and butyrate-producing functions

Dominant genera in the C-assemblage included butyrate-producing bacteria. Thus, we examined correlations between microbial assemblages and butyrate-producing functions (K00929: butyrate kinase, K01034: acetate CoA/acetoacetate CoA-transferase alpha subunit, and K01035: acetate CoA/acetoacetate CoA-transferase beta subunit). Fig. 8 indicates the Pearson’s correlation coefficients between microbial assemblages and butyrate-producing functions, showing that the C-assemblage is positively correlated with all three functions (*p* < 0.01, two-sided test, after Benjamini–Hochberg correction). However, the P- and R-assemblages were negatively correlated with some functions, and B-assemblage was positively correlated with only K00929, concurrent with the finding that *Bacteroides fragilis* has only K00929 of three functions[30].

**Figure 8:**
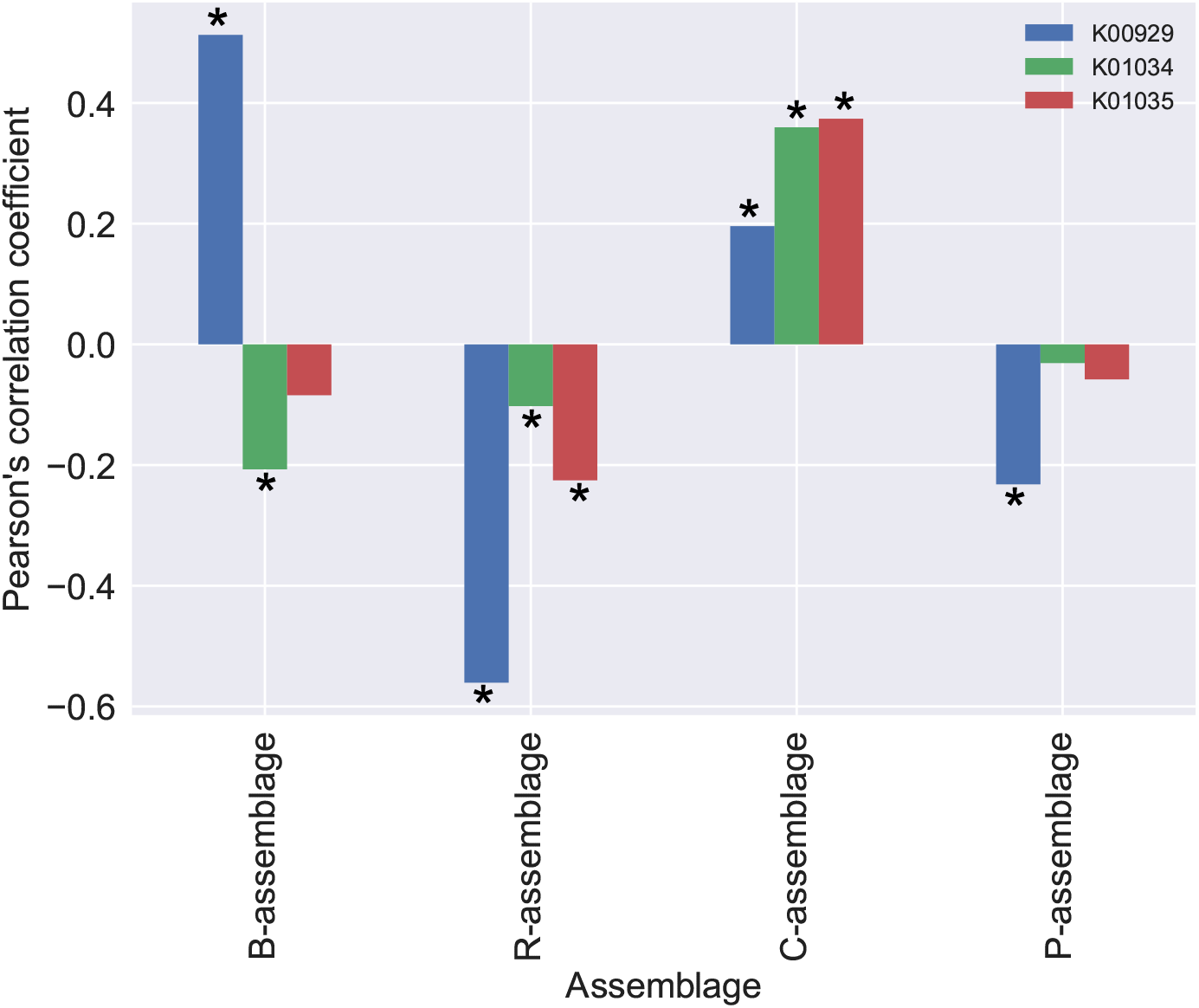
The Pearson’s correlation coefficients among the 4 assemblages and 3 butyrate-producing functions. The *x*- and *y*-axes represent the assemblages and Pearson’s correlation coefficients, respectively. Each bar of each assemblage indicates K01034, K00929, and K01035 from the left, respectively. Asterisks indicate significant differences. Significance was determined at *p* < 0.01 (two-sided test, after Benjamini–Hochberg correction).

### Functional profiles of each microbial assemblage

To discuss the functional profiles of the microbial assemblages, we applied LDA for individual functional profiles, using the same *K* number as microbial assemblages, referring to each of them as *functional assemblages*. We first investigated the correlation between microbial assemblages and functional assemblages. Supplementary Figure S6 (Additional File 1) shows that functional assemblages have a one-to-one correspondence with microbial assemblages. Therefore, we regarded functional assemblages as functional profiles of microbial assemblages in further analyses. We determined the abundances of functional categories for each assemblage (Additional File 1: Figure S7) and investigated the assemblages with the largest relative abundance for each functional category (Table 1). This table shows that metabolic functions of glycan/lipid, terpenoid/nucleotide, and vitamin/amino acid are abundant in B-, P-, and R-assemblages, respectively. In addition, no metabolic functions were abundant in the C-assemblage; however, general functional categories including the immune system and translation are abundant.

**Table 1:** The assemblage having the largest relative abundance for each functional category

| **Functional category** | **Functional assemblage** |
| --- | --- |
| Biosynthesis of Other Secondary Metabolites | B-assemblage(ko) |
| Carbohydrate Metabolism | B-assemblage(ko) |
| Digestive System | B-assemblage(ko) |
| Endocrine System | B-assemblage(ko) |
| Endocrine and Metabolic Diseases | B-assemblage(ko) |
| Environmental Adaptation | B-assemblage(ko) |
| Glycan Biosynthesis and Metabolism | B-assemblage(ko) |
| Lipid Metabolism | B-assemblage(ko) |
| Nervous System | B-assemblage(ko) |
| Transport and Catabolism | B-assemblage(ko) |
| Cell Growth and Death | P-assemblage(ko) |
| Folding, Sorting, and Degradation | P-assemblage(ko) |
| Infectious Diseases | P-assemblage(ko) |
| Metabolism of Terpenoids and Polyketides | P-assemblage(ko) |
| Nucleotide Metabolism | P-assemblage(ko) |
| Replication and Repair | P-assemblage(ko) |
| Membrane Transport | R-assemblage(ko) |
| Metabolism of Cofactors and Vitamins | R-assemblage(ko) |
| Metabolism of Other Amino Acids | R-assemblage(ko) |
| Transcription | R-assemblage(ko) |
| Xenobiotic Biodegradation and Metabolism | R-assemblage(ko) |
| Immune Diseases | R-assemblage(ko) |
| Energy Metabolism | R-assemblage(ko) |
| Amino Acid Metabolism | R-assemblage(ko) |
| Cancers | C-assemblage(ko) |
| Cell Motility | C-assemblage(ko) |
| Immune System | C-assemblage(ko) |
| Neurodegenerative Diseases | C-assemblage(ko) |
| Signal Transduction | C-assemblage(ko) |
| Translation | C-assemblage(ko) |

## Discussion

In this study, we used LDA for the detection of microbial assemblages in population-scale human gut microbiome data and discovered four microbial assemblages. Among these assemblages, while three assemblages (B-, P-, and R-assemblages) specifically emerged in the corresponding enterotypes (B-, P-, and R-types), the C-assemblage was frequently observed in every enterotype. As conventional cluster analysis of the sample focuses on the dominant genus of a cluster and the differences among clusters, the existence of non-dominant but shared microbial assemblages among individuals may have been overlooked. The detection of the C-assemblage suggested that LDA is a powerful approach for revealing the assemblage structure in massive metagenomic datasets. In addition, we found that LDA can estimate assemblages as groups of correlated microbes from the correlation analysis.

To determine *K*, i.e., the number of assemblages, we used the ad-hoc method to capture characteristics for each enterotype with high interpretability. This task, called “model selection,” is typically difficult for mixed models. Some methods for this task have been suggested [31, 32]. Yan *et al*. used cross-validation, one of the methods selecting the model with the highest likelihood against the test data. However, these methods tend to overestimate *K*, leading to presenting difficulties in clarifying the association between enterotypes and assemblages. Indeed, Yan *et al*. estimated *K* = 60, although the number of samples was less than that of the samples used in this study.

As mentioned above, the genera mainly appearing in the B- and P-assemblages tend to occur in the B- and P-types, respectively. The genera specifically appearing in the B- and P-types were reported to have functions for metabolizing protein/animal fat and carbohydrates, respectively [33], and the genera mainly appearing in the B- and P-assemblages may consequently have functions for metabolizing protein/animal fat and carbohydrates, respectively. We could confirm that lipid metabolism functions were abundant in the B-assemblage through functional assemblage analysis. This result suggests that the B-assemblage in the human gut becomes dominant through a fat-rich diet. In the same way, the genera mainly appearing in the C-assemblage may have functions that do not correspond with dietary habits because they appeared in all enterotypes. This suggestion is concurrent with the finding that immune cells and translation are abundant in the C-assemblage.

There are two notable issues about the C-assemblage. First, the C-assemblage can coexist with all of the other three assemblages, which was found in almost all countries. In other words, genera mainly appearing in the C-assemblage were generalists in the human gut environment [34, 35]. While generalists can adapt to diverse environments, they were not specialized to particular environments unlike specialists. This difference in survival strategy may be the reason the genera mainly appearing in the C-assemblage were not dominant genera in the human gut microbiome. Second, it is therefore possible that the C-assemblage is the core gut microbiome [9, 36]. However, C-assemblage abundance is not consistent from person to person; as such, what determines the existence of C-assemblage in the gut microbiome? The dominant genera of the C-assemblage (*i.e., Clostridium, Eubacterium, Faecalibacterium*, *Roseburia*, *Coprococcus*, and *Butyrivibrio*) include representative butyrate-producing species (Table 2) [37, 38]. In addition, we found that the C-assemblage had correlations with the three butyrate-producing functions. Butyrate is known to have anti-inflammatory effects [39] and to be associated with IBD, type-2 diabetes, and colorectal cancer [40, 41, 42]. Therefore, C-assemblage abundance may indicate the health of hosts, although the dataset used in this study contains only healthy individuals. In addition, we found that ages and BMI did not relate to the presence of the C-assemblage (Additional File 1: Figure S8). Further research is accordingly required, such as comparisons of C-assemblage abundance between individuals with and without a disease.

**Table 2:** Dominant genera of the C-assemblage and the probability of the C-assemblage as estimated by LDA.

| **Genus** | **Probability** |
| --- | --- |
| *Clostridium* | 0.179865 |
| *Eubacterium* | 0.150802 |
| *Unclassified Firmicutes* | 0.129783 |
| *Faecalibacterium* | 0.093720 |
| *Ruminococcus* | 0.085272 |
| *Roseburia* | 0.074214 |
| *Alistipes* | 0.072359 |
| *Coprococcus* | 0.029497 |
| *Butyrivibrio* | 0.021738 |

We envision two future directions for applications of LDA to metagenomic data. The first is the application to more diverse datasets. Metagenomic data have been sampled from not only human guts but also various environments such as the atmosphere [43], ocean [44], and soil [45]. Application of LDA to these data should help reveal the structure of microbial assemblages at a global-scale [46]. The second is the extension of the LDA model because LDA has high model extensibility. Indeed, many extended LDA models have been proposed for natural language processing [47, 48, 49, 50]. The application of these extended LDA models to metagenomic analysis is a fascinating research focus for further elucidation of microbial assemblage structure. For example, applying supervised topic models [51], which utilize label information to estimate assemblage structures, to patient metagenomic data could detect microbial assemblages related to disease. The pachinko allocation model [47], which models hierarchical assemblage structures, may be useful for revealing sub-assemblages within an assemblage. A transition in assemblage composition could be estimated from time-series data from human gut microbiomes[52] using the topic tracking model[50].

## Conclusions

In this study, we conducted an assemblage analysis on a large-scale human gut metagenome dataset using LDA. While three assemblages specifically emerged that corresponded to enterotypes, the C-assemblage was frequently observed in all three enterotypes. Interestingly, the dominant genera of the C-assemblage include representative butyrate-producing species. LDA is a powerful method for detecting microbial assemblages, and it has the potential to reveal the structure of the human gut microbiome.

## Supporting information

This file includes Figures S1, S2, S3, S4, S5, S6, S7, S8, and Table S1.

## List of abbreviations

LDA: latent Dirichlet allocation
IBD: inflammatory bowel disease
PAM: partitioning around medoids
DMM: Dirichlet multinomial mixture
VB: variational Bayes
VLB: variational lower-bound

## Ethics approval and consent to participate

Not applicable.

## Consent for publication

Not applicable.

## Availability of data and material

Supplementary material is available from the journal website.

## Funding

This work was supported by the Ministry of Education, Culture, Sports, Science and Technology (KAKENHI) (grant numbers JP16H05879, JP16H01318, JP16H02484, and 17K20032 to MH)

## Competing interests

The authors declare that they have no competing interests.

## Author’s contributions

M. Hamada and TF conceived the study. M. Hamada supervised this study. SN and M. Hattori processed the data. SH implemented the method and performed all the computational experiments. SH, SN, TF, and M. Hamada analyzed the results. TF, SH, and M. Hamada wrote the draft manuscript, and SN and M. Hattori revised it critically. All authors read and approved the final manuscript.

## Acknowledgements

The computations in this research were performed using the supercomputing facilities at the National Institute of Genetics in Research Organization of Information and Systems.

## Additional Files

**Additional File 1 — supplementary.pdf**

This file includes Figures S1, S2, S3, S4, S5, S6, S7, S8, and Table S1.

