## Supplementary material for "Revealing microbial assemblage structure in the human gut microbiome using latent Dirichlet allocation": This file includes Figures S1, S2, S3, S4, S5, S6, S7, S8, and Table S1.

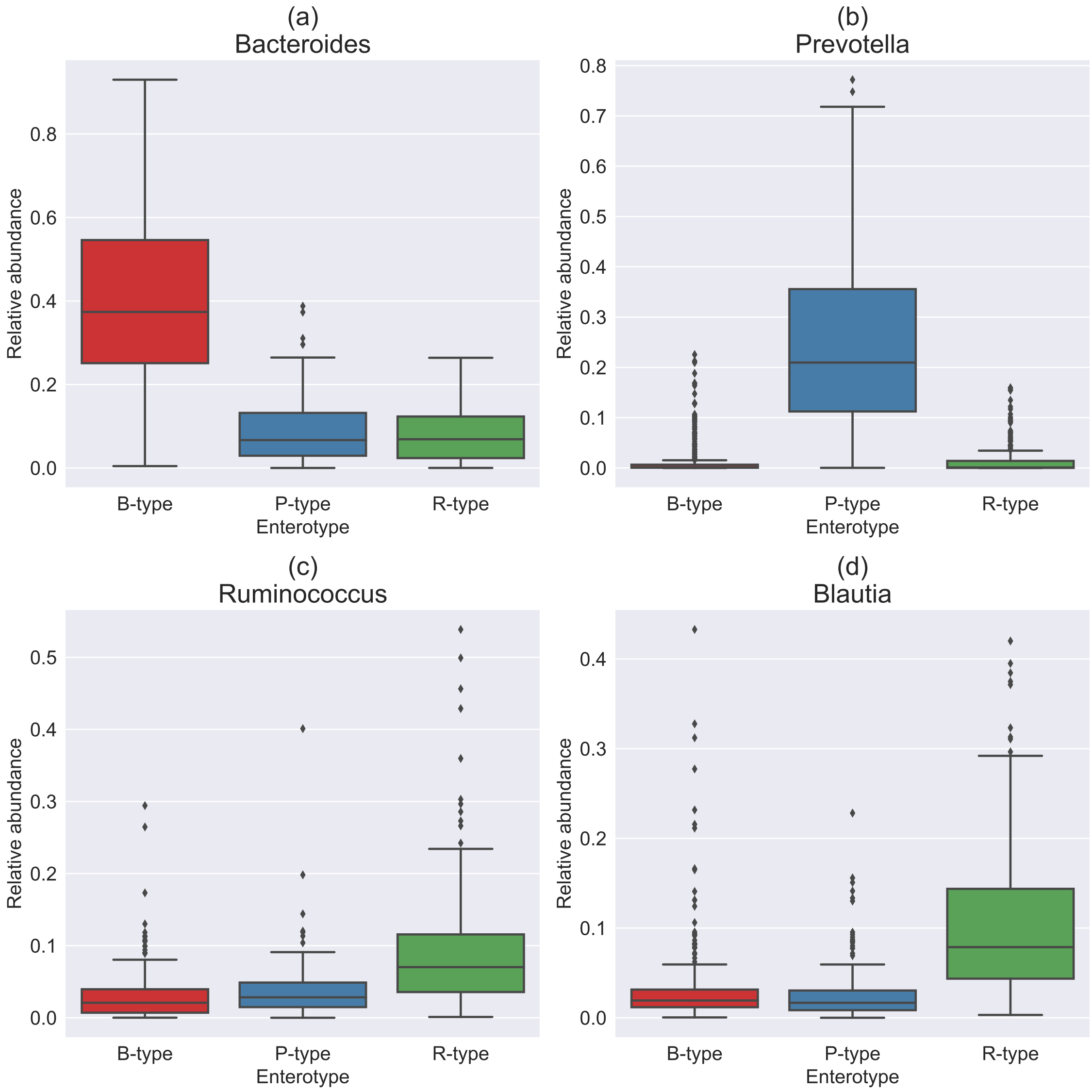


Figure S1: The relative abundance of 4 major genera ((a)*Bacteroides*, (b)*Prevotella*, (c)*Ruminococcus*, (d)*Blautia*) for each enterotype. The $x$- and $y$-axes represent enterotype and relative abundance, respectively.

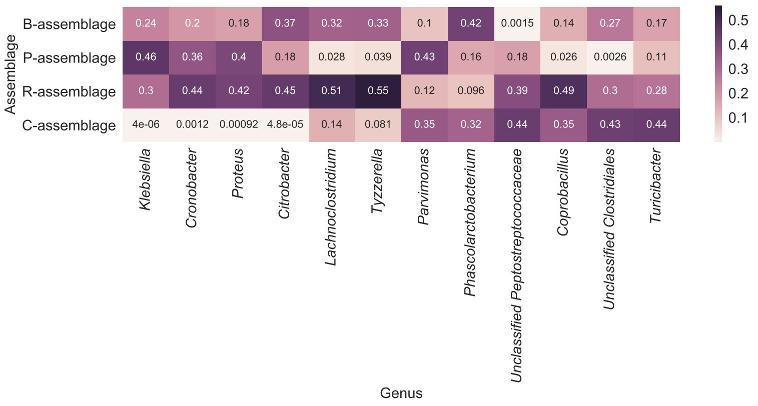


Figure S2: The $P(a_k|g_j)$ (Eq. 3 in the main text) of the genera with hith entropy score. The $x$- and $y$-axes represent genus and microbial assemblage, respectively.

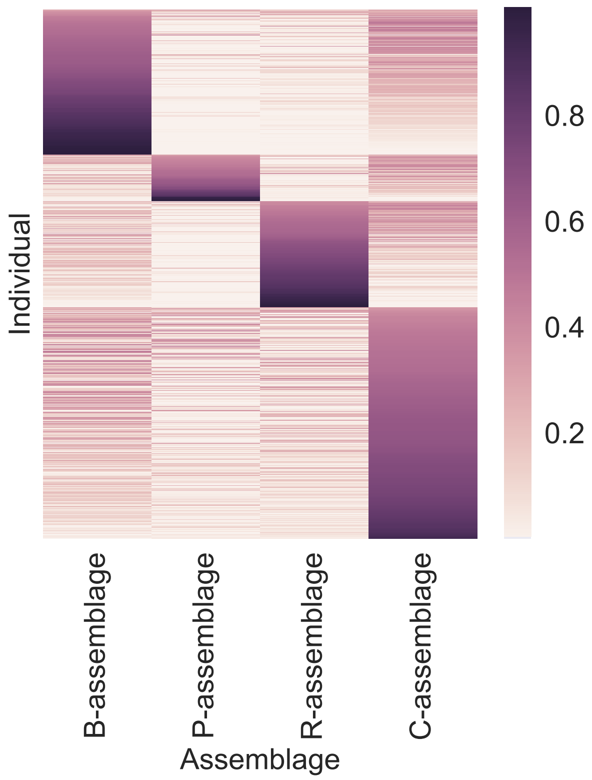


Figure S3: Estimated microbial assemblage distribution for each individual. The $x$- and $y$-axes represent microbial assemblages and individual samples, respectively. Individuals are ordered by the probability of the dominant assemblage.

Table S1: The entropy score of $P(a_k|g_j)$ for all genera used.

| Genus | entropy |
|---|---|
| *Turicibacter* | 1.260681 |
| *Phascolarctobacterium* | 1.245466 |
| *Parvimonas* | 1.220488 |
| *Unclassified Clostridiales* | 1.092140 |
| *Coprobacillus* | 1.088414 |
| *Lachnoclostridium* | 1.082888 |
| *Klebsiella* | 1.061892 |
| *Cronobacter* | 1.056325 |

Table S1: The entropy score of $P(a_k|g_j)$ for all genera used. – *Continued from previous page*

| Genus | entropy |
|---|---|
| *Unclassified Peptostreptococcaceae* | 1.044608 |
| *Proteus* | 1.042833 |
| *Citrobacter* | 1.037434 |
| *Tyzzerella* | 1.021686 |
| *Veillonella* | 0.999699 |
| *Holdemania* | 0.983926 |
| *Yersinia* | 0.980842 |
| *Bilophila* | 0.951916 |
| *Faecalibacterium* | 0.937553 |
| *Unclassified Lachnospiraceae* | 0.933686 |
| *Paraprevotella* | 0.926816 |
| *Enterobacter* | 0.924527 |
| *Unclassified Bacteria* | 0.906336 |
| *Subdoligranulum* | 0.893402 |
| *Pseudomonas* | 0.891144 |
| *Peptoclostridium* | 0.880690 |
| *Flavonifractor* | 0.878429 |
| *Eubacterium* | 0.857729 |
| *Lachnospira* | 0.840097 |
| *Dorea* | 0.822750 |
| *Unclassified Erysipelotrichaceae* | 0.816711 |
| *Escherichia* | 0.814104 |
| *Megasphaera* | 0.812753 |
| *Roseburia* | 0.788690 |
| *Erysipelatoclostridium* | 0.784843 |
| *Acidaminococcus* | 0.773040 |
| *Dielma* | 0.756564 |
| *Intestinibacter* | 0.744818 |
| *Mycobacterium* | 0.736164 |
| *Sutterella* | 0.725791 |
| *Scardovia* | 0.718660 |
| *Olsenella* | 0.713303 |
| *Providencia* | 0.709675 |
| *Stomatobaculum* | 0.709085 |
| *Odoribacter* | 0.704806 |
| *Unclassified Clostridiales Family XIII. Incerta...* | 0.703168 |
| *Peptostreptococcus* | 0.697437 |
| *Kandleria* | 0.696656 |
| *Marvinbryantia* | 0.695373 |
| *Cloacibacillus* | 0.695198 |
| *Sharpea* | 0.695112 |
| *Atopobium* | 0.694374 |
| *Barnesiella* | 0.693038 |
| *Catenibacterium* | 0.687024 |
| *Tannerella* | 0.686389 |
| *Streptococcus* | 0.685115 |
| *Oenococcus* | 0.684101 |
| *Unclassified Bacteroidales* | 0.683936 |
| *Dialister* | 0.683460 |
| *Azospirillum* | 0.680177 |
| *Weissella* | 0.677471 |
| *Ruminococcus* | 0.673173 |
| *Megamonas* | 0.669763 |
| *Leuconostoc* | 0.668794 |
| *Unclassified Burkholderiales* | 0.665163 |

Table S1: The entropy score of $P(a_k|g_j)$ for all genera used. – *Continued from previous page*

| Genus | entropy |
|---|---|
| *Methanosphaera* | 0.654763 |
| *Blautia* | 0.646975 |
| *Solobacterium* | 0.644240 |
| *Enterococcus* | 0.644030 |
| *Akkermansia* | 0.631433 |
| *Alistipes* | 0.624308 |
| *Actinomyces* | 0.622316 |
| *Unclassified Ruminococcaceae* | 0.620387 |
| *Peptoniphilus* | 0.616613 |
| *Senegalimassilia* | 0.595117 |
| *Pseudoflavonifractor* | 0.594075 |
| *Unclassified Proteobacteria* | 0.591897 |
| *Haemophilus* | 0.588196 |
| *Desulfovibrio* | 0.585395 |
| *Alloprevotella* | 0.582315 |
| *Actinobaculum* | 0.579283 |
| *Propionibacterium* | 0.578085 |
| *Oxalobacter* | 0.576018 |
| *Mitsuokella* | 0.571258 |
| *Coprococcus* | 0.562809 |
| *Fusobacterium* | 0.561115 |
| *Adlercreutzia* | 0.546762 |
| *Butyricicoccus* | 0.543292 |
| *Anaerostipes* | 0.539088 |
| *Thauera* | 0.535747 |
| *Pedobacter* | 0.535627 |
| *Luteimonas* | 0.535625 |
| *Allobaculum* | 0.535624 |
| *Sphaerochaeta* | 0.535624 |
| *Kallipyga* | 0.535624 |
| *Lamprocystis* | 0.535565 |
| *Sphingobacterium* | 0.535565 |
| *Sphingopyxis* | 0.535565 |
| *Unclassified Aminicenantes* | 0.535499 |
| *Brevibacillus* | 0.535499 |
| *Campylobacter* | 0.521678 |
| *Methanobrevibacter* | 0.520127 |
| *Cryptobacterium* | 0.513993 |
| *Ruminiclostridium* | 0.510680 |
| *Eggerthella* | 0.509646 |
| *Lactococcus* | 0.504261 |
| *Butyricimonas* | 0.491595 |
| *Unclassified Bacteroidetes* | 0.474846 |
| *Shewanella* | 0.459807 |
| *Porphyromonas* | 0.434124 |
| *Parabacteroides* | 0.427271 |
| *Johnsonella* | 0.423664 |
| *Corynebacterium* | 0.420475 |
| *Brachyspira* | 0.393297 |
| *Gemella* | 0.377228 |
| *Gordonibacter* | 0.369317 |
| *Lactobacillus* | 0.347828 |
| *Unclassified Xanthomonadaceae* | 0.339638 |
| *Dickeya* | 0.339637 |
| *Kosakonia* | 0.339551 |

Table S1: The entropy score of $P(a_k|g_j)$ for all genera used. – *Continued from previous page*

| Genus | entropy |
|---|---|
| *Moraxella* | 0.339550 |
| *Arthrospira* | 0.339549 |
| *Anaeroglobus* | 0.339549 |
| *Carnobacterium* | 0.339549 |
| *Shinella* | 0.339507 |
| *Stenotrophomonas* | 0.339459 |
| *Allofustis* | 0.339458 |
| *Lelliottia* | 0.339458 |
| *Bacillus ¡bacterium¿* | 0.336655 |
| *Hafnia* | 0.325478 |
| *Granulicatella* | 0.298182 |
| *Collinsella* | 0.294826 |
| *Actinobacillus* | 0.253157 |
| *Mobiluncus* | 0.253157 |
| *Cupriavidus* | 0.253089 |
| *Unclassified Propionibacteriaceae* | 0.253089 |
| *Shuttleworthia* | 0.253055 |
| *Ethanoligenens* | 0.253055 |
| *Finegoldia* | 0.253018 |
| *Shigella* | 0.252294 |
| *Coprobacter* | 0.251584 |
| *Rikenella* | 0.240003 |
| *Parasutterella* | 0.239172 |
| *Lachnoanaerobaculum* | 0.235800 |
| *Clostridium* | 0.216568 |
| *Sinorhizobium* | 0.203630 |
| *Aeromicrobium* | 0.203630 |
| *Tetrasphaera* | 0.203574 |
| *Janthinobacterium* | 0.203572 |
| *Dysgonomonas* | 0.203572 |
| *Microlunatus* | 0.203546 |
| *Pantoea* | 0.203546 |
| *Synergistes* | 0.203514 |
| *Unclassified Firmicutes* | 0.174778 |
| *Vibrio* | 0.171243 |
| *Anaerovibrio* | 0.171171 |
| *Bulleidia* | 0.171171 |
| *Unclassified Oxalobacteraceae* | 0.171171 |
| *Lachnobacterium* | 0.171144 |
| *Raoultella* | 0.169913 |
| *Abiotrophia* | 0.169097 |
| *Anaerotruncus* | 0.168921 |
| *Streptomyces* | 0.157833 |
| *Thermus* | 0.148239 |
| *Trueperella* | 0.148238 |
| *Serratia* | 0.148238 |
| *Macrococcus* | 0.148238 |
| *Terrisporobacter* | 0.148238 |
| *Parascardovia* | 0.148238 |
| *Paenibacillus* | 0.136984 |
| *Riemerella* | 0.131086 |
| *Sphingomonas* | 0.131048 |
| *Candidatus Arthromitus* | 0.131048 |
| *Negativicoccus* | 0.131030 |
| *Bacteroides* | 0.122743 |

Table S1: The entropy score of $P(a_k|g_j)$ for all genera used. – *Continued from previous page*

| Genus | entropy |
|---|---|
| *Candidatus Stoquefichus* | 0.117656 |
| *Aggregatibacter* | 0.112499 |
| *Pseudoalteromonas* | 0.106937 |
| *Unclassified Betaproteobacteria* | 0.106937 |
| *Robinsoniella* | 0.106905 |
| *Filifactor* | 0.106891 |
| *Brevibacterium* | 0.106891 |
| *Kocuria* | 0.098099 |
| *Unclassified Alphaproteobacteria* | 0.098099 |
| *Georgenia* | 0.098099 |
| *Mogibacterium* | 0.098071 |
| *Paracoccus* | 0.098056 |
| *Morganella* | 0.090671 |
| *Rothia* | 0.089105 |
| *Nocardioides* | 0.084365 |
| *Staphylococcus* | 0.082676 |
| *Unclassified Coriobacteriaceae* | 0.078953 |
| *Anaerofustis* | 0.078953 |
| *Slackia* | 0.078174 |
| *Pasteurella* | 0.070096 |
| *Aerococcus* | 0.070047 |
| *Anaerococcus* | 0.066380 |
| *Anaerosalibacter* | 0.057431 |
| *Unclassified Candidatus Saccharibacteria* | 0.057431 |
| *Gardnerella* | 0.052776 |
| *Plesiomonas* | 0.050727 |
| *Xanthomonas* | 0.048869 |
| *Elizabethkingia* | 0.048838 |
| *Succinimonas* | 0.047110 |
| *Selenomonas* | 0.046316 |
| *Oribacterium* | 0.044003 |
| *Beggiatoa* | 0.041308 |
| *Unclassified Bacteroidaceae* | 0.040074 |
| *Acidovorax* | 0.035891 |
| *Bifidobacterium* | 0.035527 |
| *Micrococcus* | 0.029757 |
| *Catonella* | 0.029757 |
| *Aeromonas* | 0.028547 |
| *Leclercia* | 0.026936 |
| *Kluyvera* | 0.026434 |
| *Burkholderia* | 0.025968 |
| *Geobacillus* | 0.025960 |
| *Alloscardovia* | 0.025499 |
| *Rhodococcus* | 0.025052 |
| *Prevotella* | 0.024904 |
| *Microbacterium* | 0.024628 |
| *Acinetobacter* | 0.023983 |
| *Varibaculum* | 0.023434 |
| *Listeria* | 0.022350 |
| *Alcanivorax* | 0.022160 |
| *Eikenella* | 0.020773 |
| *Oscillibacter* | 0.020362 |
| *Enterorhabdus* | 0.018686 |
| *Helicobacter* | 0.017187 |
| *Bifidobacterium* | 0.013555 |

Table S1: The entropy score of $P(a_k|g_j)$ for all genera used. – *Continued from previous page*

| Genus | entropy |
|---|---|
| *Methylobacterium* | 0.012762 |
| *Leptotrichia* | 0.012559 |
| *Pyramidobacter* | 0.010146 |
| *Lysinibacillus* | 0.009544 |
| *Kingella* | 0.008453 |
| *Bordetella* | 0.007775 |
| *Cardiobacterium* | 0.007596 |
| *Methanomassiliicoccus* | 0.006476 |
| *Lautropia* | 0.005972 |
| *Succinatimonas* | 0.004608 |
| *Spiroplasma* | 0.004588 |
| *Acidiphilium* | 0.003029 |
| *Corallococcus* | 0.002747 |
| *Achromobacter* | 0.002647 |
| *Capnocytophaga* | 0.002549 |
| *Cetobacterium* | 0.002347 |
| *Butyrivibrio* | 0.002281 |
| *Coraliomargarita* | 0.002105 |
| *Enorma* | 0.001869 |
| *Acholeplasma* | 0.000901 |
| *Pediococcus* | 0.000880 |
| *Neisseria* | 0.000365 |
| *Treponema* | 0.000171 |
| *Acetobacter* | 0.000142 |

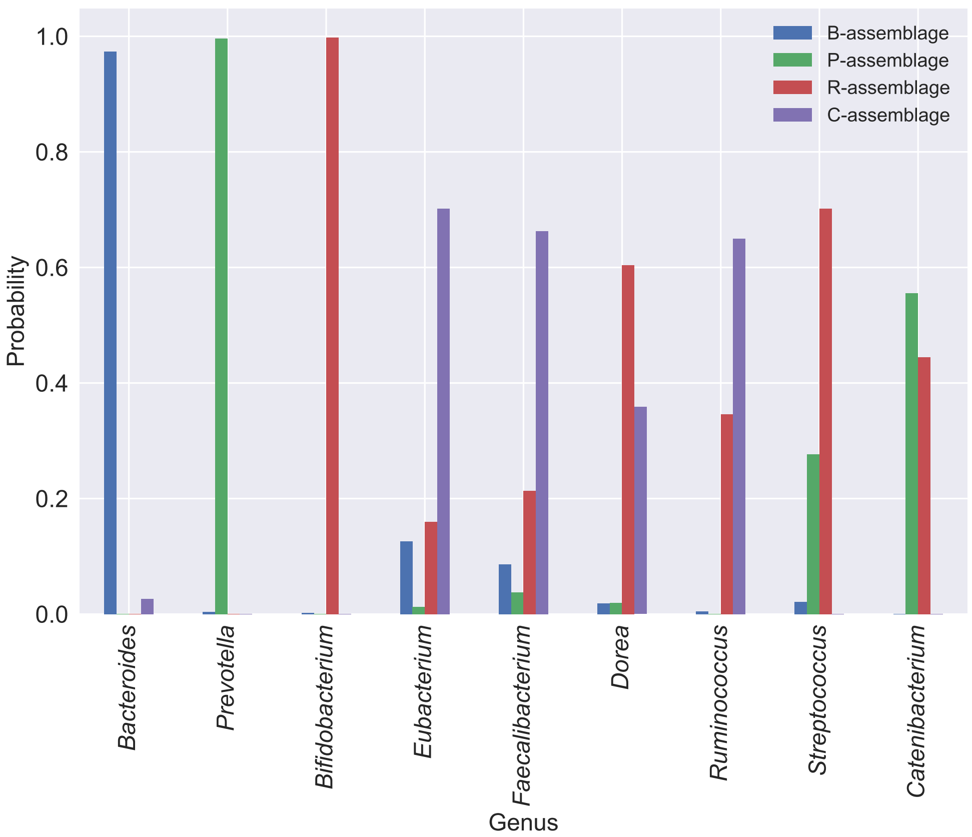


Figure S4: The $P(a_k|g_j)$ of *Bacteroides*, *Prevotella*, *Bifidobacterium* and the genera that correlate with the other assemblage genera. The $x$- and $y$-axes represent the genus and the $P(a_k|g_j)$, respectively.

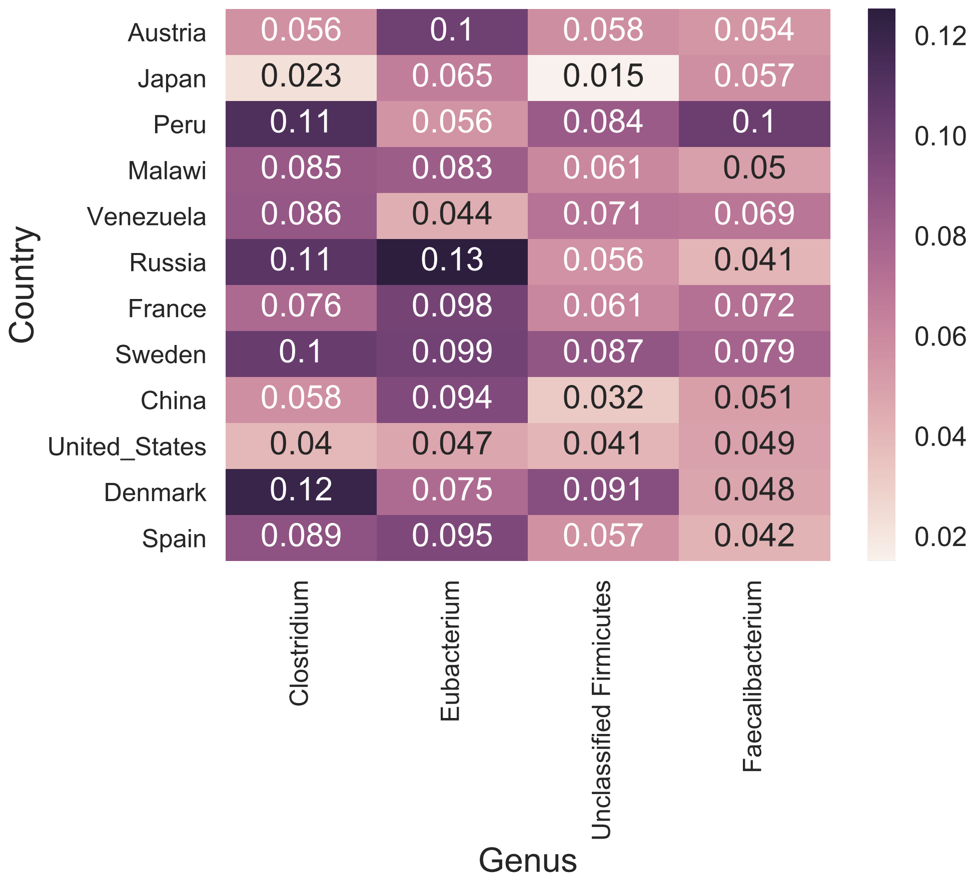


Figure S5: The average relative abundance of *Clostridium*, *Eubacterium*, Unclassified *Firmicutes*, and *Faecalibacterium*, for each country. The $x$- and $y$-axes represent the genus and the country of the individual, respectively.

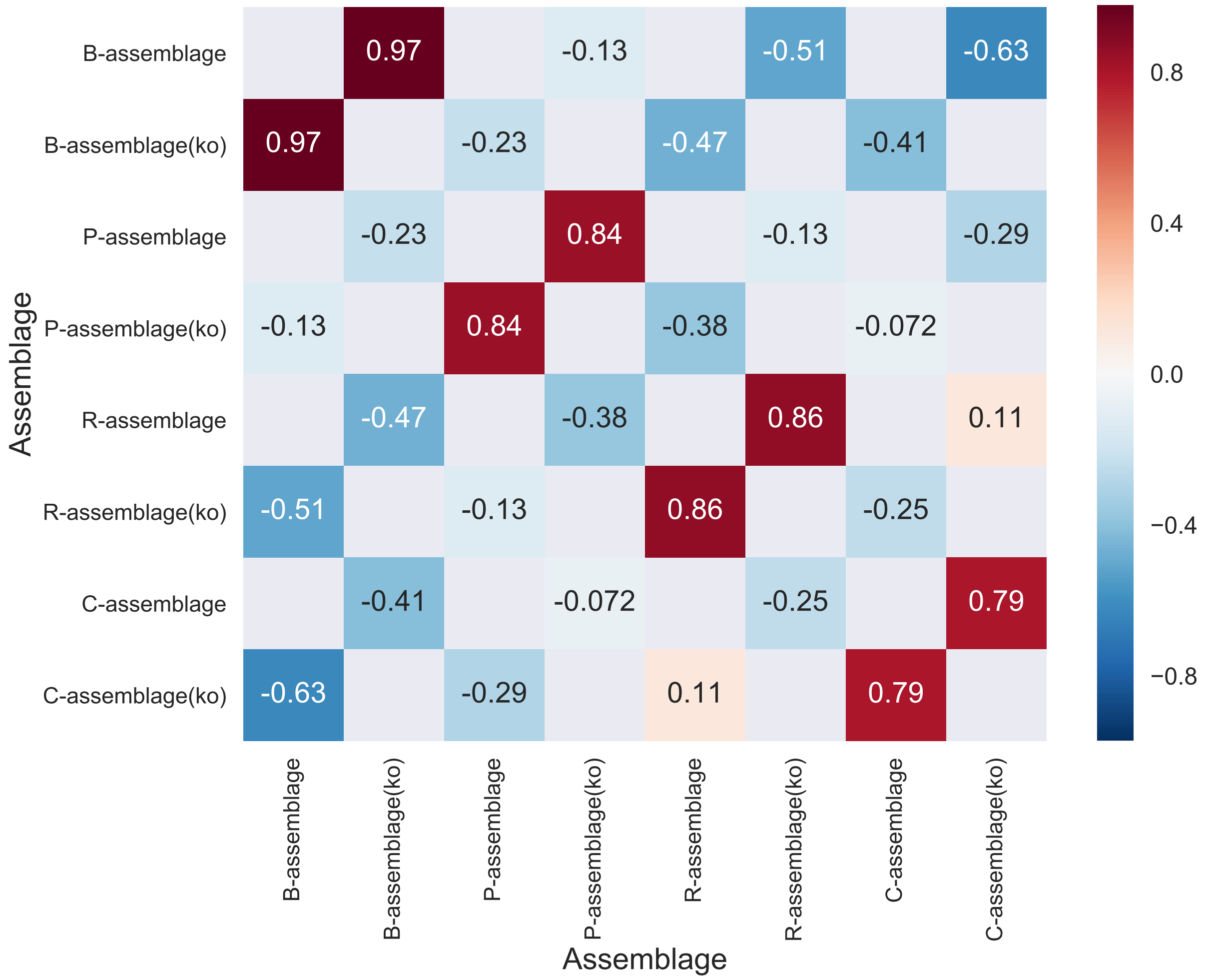


Figure S6: Pearson's correlation coefficients among microbial assemblages and functional assemblages. Both of the $x$- and $y$-axes represent assemblages.

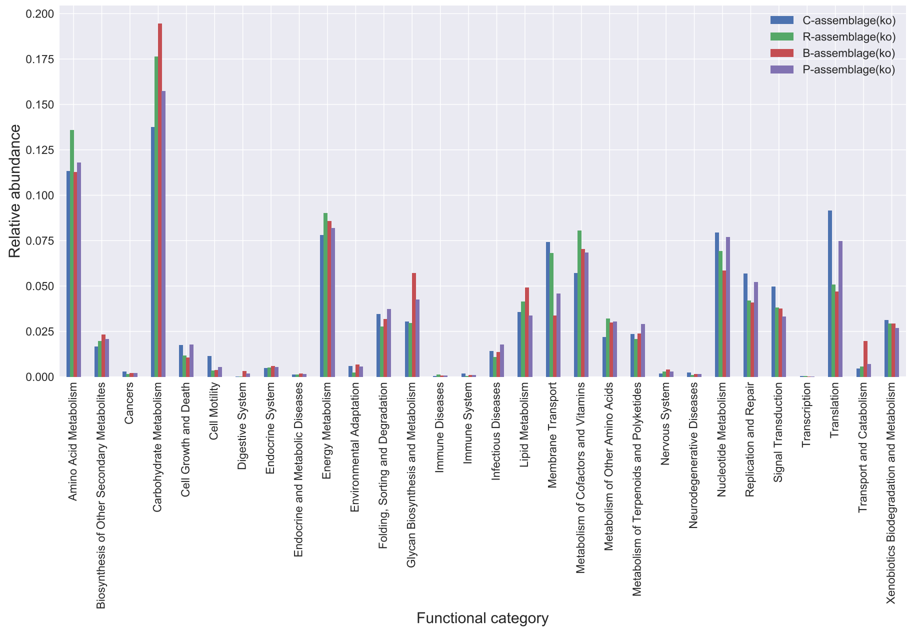


Figure S7: The relative abundances of functional categories for each assemblage. The $x$- and $y$-axes represent the function category and the relative abundance, respectively.

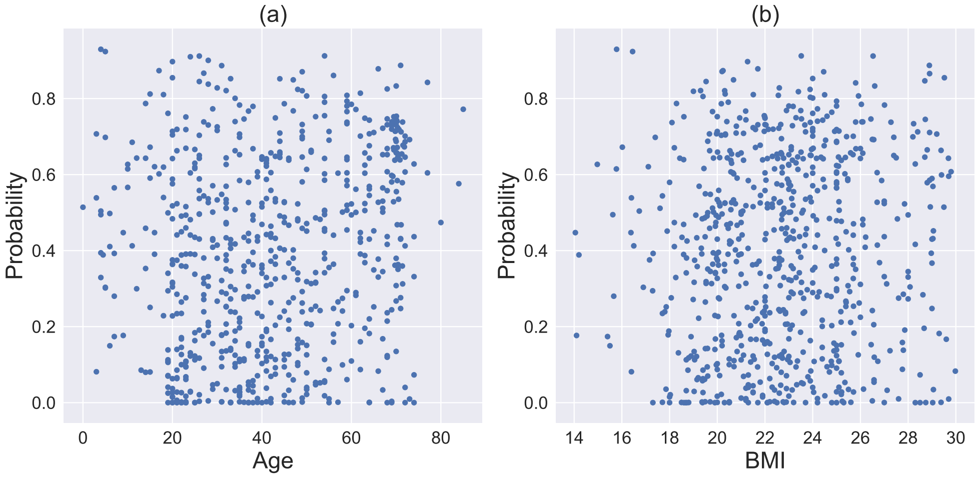


Figure S8: The relationship between C-assemblage probability and two metadata ((a)age and (b)BMI). The $x$- and $y$-axes represent metadata and C-assemblage probability, respectively.
